# Recurrent interactions between olfactory bulb and olfactory cortex support cross-sniff perceptual continuity in humans

**DOI:** 10.64898/2026.08.31.747807

**Authors:** Frans Nordén, Anja L. Winter, Leslie M. Kay, Artin Arshamian, Mikael Lundqvist, Johan N. Lundström

## Abstract

Olfactory perception relies on active sampling, with successive inhalations providing discrete sensory inputs that in humans can be separated by several seconds. Yet odors are perceived as continuous and stable, raising the question of how the brain integrates sensory information across these temporal gaps. Here, we used electrobulbogram (EBG) recordings in 48 participants to demonstrate that successive sniffs are not processed independently but are linked through recurrent oscillatory dynamics between the olfactory bulb (OB) and piriform cortex (PC). Specifically, first-sniff alpha/beta activity in the OB and OB–PC functional connectivity predicted second-sniff gamma power, while transfer entropy indicated a directional temporal dependence from first-sniff alpha/beta to second-sniff gamma dynamics. In parallel, second-sniff gamma activity selectively tracked subjective odor valence prior to inhalation onset. At the network level, the PC exhibited stronger alpha/beta-band connectivity with orbitofrontal, insular, and prefrontal regions during the first sniff than during the second, suggesting that early evaluative processing provides a contextual signal that is carried forward to shape subsequent sensory representations. These results demonstrate that the human OB-PC circuit carries evaluative information across inhalations through directed alpha/beta-to-gamma interactions, providing a mechanism for maintaining perceptual continuity in a sensory system defined by temporally discrete sampling.

## Introduction

Olfactory perception relies on active sampling where each inhalation draws a bolus of odorant-filled air, generating a discrete snapshot of the chemical environment. Unlike vision and audition, where the sensory surfaces are continuously stimulated, olfactory input arrives in temporally separated packets bound to the respiratory cycle. In humans, the olfactory percept often needs to be preserved in the absence of sensory input for several seconds, yet it remains stable. How the olfactory system integrates information across these successive samples of an ongoing odor to construct a stable percept remains a central, yet largely unresolved, question in human olfactory neuroscience.

In rodents, where much of the foundational work has been conducted, the oscillatory architecture of the olfactory bulb (OB) and piriform cortex (PC) provides a candidate mechanism for such integration. Odor presentation engages distinct frequency bands in a coordinated manner: Gamma-band activity (30–100 Hz) is expressed during early sensory processing and is closely linked to OB-PC interactions, whereas beta-band activity (12–30 Hz) emerges during recurrent interactions involving the PC and broader cortical networks (Bressler & Freeman, 1980; Kay & Stopfer, 2006; Martin & Ravel, 2014). These dynamic interactions unfold rapidly across the fast, theta-locked sniff cycles characteristic of rodent respiration, providing a temporal scaffold that coordinates gamma- and beta-band dynamics within and across inhalations (Fontanini & Bower, 2006; Laurent et al., 2001). Within this framework, the OB is not merely a relay station but a site of active organization and reorganization, where centrifugal beta-band interactions can shape local sensory processing and gamma-band dynamics (Kay & Stopfer, 2006; Martin & Ravel, 2014).

Whether this oscillatory architecture generalizes to humans has only recently become assessable. Intracranial recordings from the human PC demonstrate that theta-band activity carries odor identity information and is phase-locked to respiration (Jiang et al., 2017), while beta and gamma oscillations support accurate odor identification (Yang et al., 2022). Recent direct recordings from the human OB have demonstrated theta-band activity that are phase-locked to respiration (Sheriff et al., 2026) and noninvasive electrobulbogram (EBG) recordings have shown that the human OB encodes odor valence in beta and gamma bands (Iravani et al., 2021; Li et al., 2026), transmits odor valence information to the PC via gamma oscillations, and receives top-down beta-band feedback (Nordén et al., 2024, 2026). If these interactions contribute to maintaining perceptual continuity across successive samples, oscillatory activity during one sniff should be associated with sensory processing during the next. However, these studies share a tacit assumption: that each sniff constitutes an independent sample, an assumption built into paradigms of brief, isolated odor presentations (typically 1–2 s) inherited from rodent work. Because rodents sniff at high frequencies, often within the theta range, a 1 s stimulus spans multiple inhalation–exhalation cycles and cortical dynamics evolve in a stereotyped manner across sniffs (Frederick et al., 2016), but in humans, who rarely exceed one sniff per second, the same duration typically captures only a single inhalation. Consequently, the inter-sniff oscillatory dynamics, central to maintaining a stable odor perception, have in humans been systematically overlooked.

If recurrent interactions between the OB and PC contribute to maintaining perceptual continuity, neural activity during one sniff should influence processing during the next. Specifically, alpha/beta activity during the first sniff may shape subsequent gamma-band responses in the OB, reflecting the impact of prior sampling history on later sensory processing. We therefore hypothesize that alpha/beta-band activity in the OB during the first sniff predicts gamma-band activity during the second sniff, and that this relationship is mediated by alpha/beta-band functional connectivity between the OB and PC. We further hypothesized that the first sniff would be accompanied by stronger functional connectivity between the PC and higher-order evaluative and associative regions. Such a pattern would be consistent with the rapid integration of sensory signals with affective and contextual processing during initial odor sampling. We tested these predictions using EBG recordings in 48 participants who rated the valence and intensity of odors across two consecutive sniffs of an ongoing odor (Figure 1A–D).

**Figure 1:**
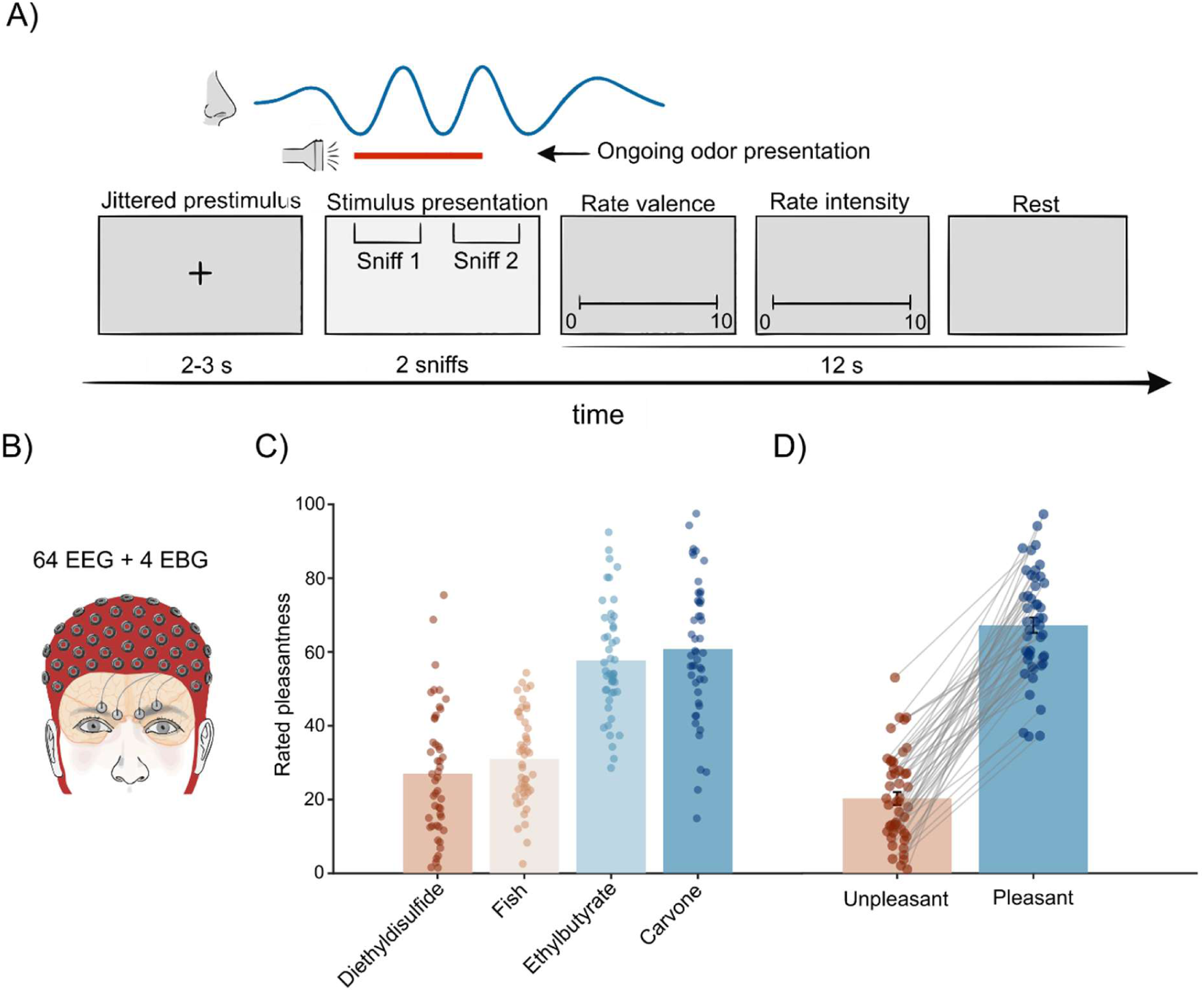
Methodology and subjective odor valence ratings for the odors. **A)** Trial overview where the blue line indicates the breathing cycle during the two sniffs during which the stimulus is presented. **B)** Illustration of the layout of the EEG and electrobulbogram electrodes. **C)** Rated valence for the four odors included in the experiment, where dots represent individual participant’s means. The color scale from terracotta to blue indicates a scale from unpleasant to pleasant. **D)** A median split of the four odors based on rated valence visualizes a distinct difference between unpleasant and pleasant odors.

## Results

### Second-sniff gamma activity encodes odor valence and emerges prior to inhalation

To determine whether successive sniffs are accompanied by distinct patterns of neural activity, we compared time-frequency responses in the olfactory bulb (OB) and piriform cortex (PC) between the first and second inhalation (Fig. 1A, Fig. 2A-D).

**Figure 2:**
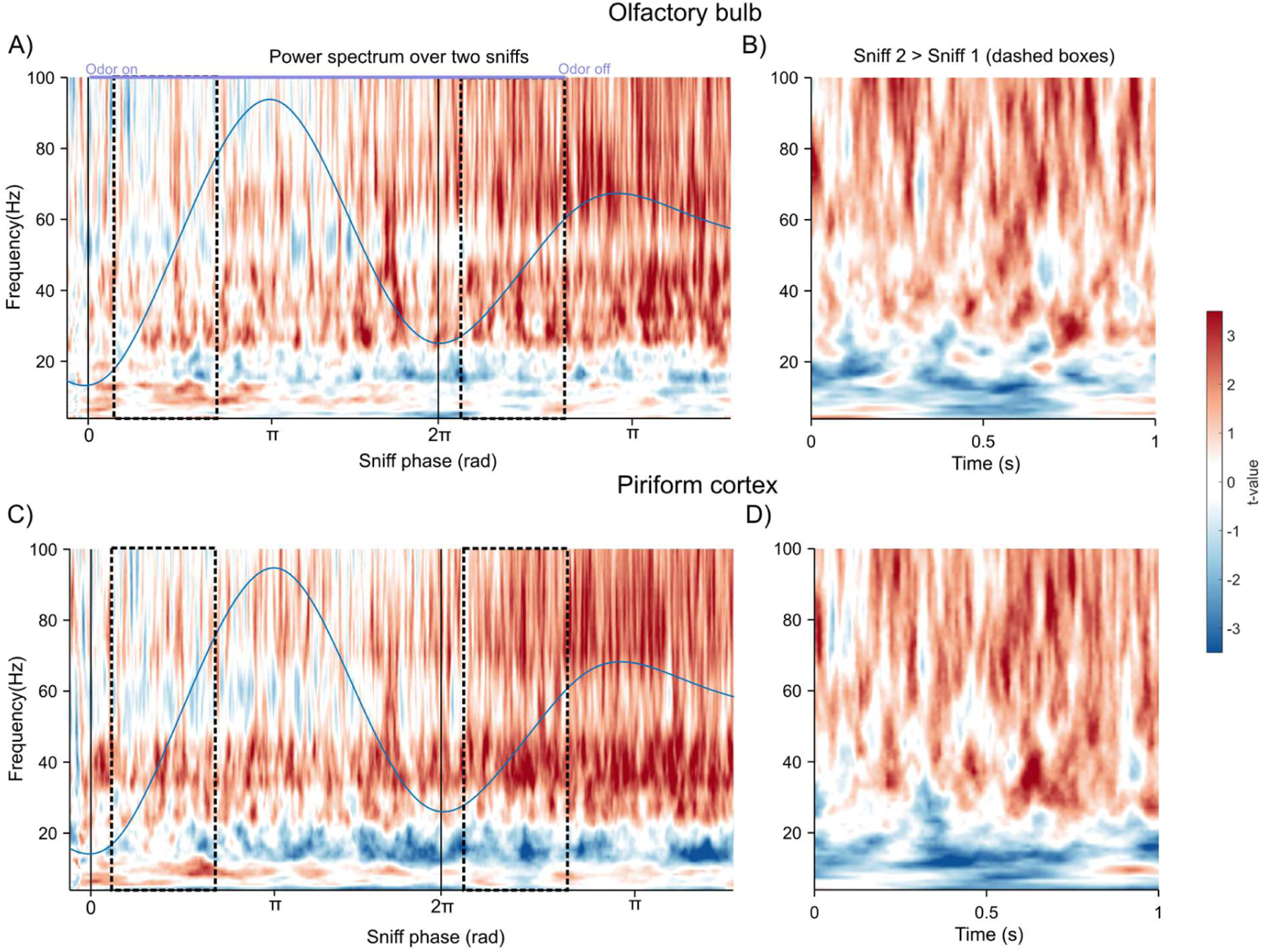
Time frequency response over two sniffs. While theta-alpha/beta activity is more prominent in the first sniff for both OB and PC, gamma activity becomes stronger over time and is stronger during the second sniff. **A)** Average olfactory bulb activity over the whole sniff cycle normalized in the time dimension. **B)** OB power spectrum where the initial second of the second and first sniff are contrasted (seen in the dashed box in **A**). **C)** Same as **A,** but for the piriform cortex. **D)** Same as **B,** but for the piriform cortex.

Both regions exhibited the canonical olfactory response, consisting of an early increase in gamma- and theta-band activity followed by later alpha/beta oscillations. Critically, the magnitude of these components differed across sniffs: theta, alpha, and beta activity were stronger during the first sniff, whereas gamma activity was significantly enhanced during the second. Cluster-corrected direct contrasts confirmed this pattern in both regions (OB: theta/alpha/beta: t = 3.9, p = .031, CI = [.021, .024]; gamma: t =−3.6, p = .026, CI = [−.025, −.019]; PC: theta/alpha/beta: t = 4.3, p = .017, CI = [0.0266, 0.0314]; gamma: t =−4.6, p = .023, CI = [−.023, −.017]), indicating a shift from low-frequency to high-frequency dominance across successive samples.

Because sampling behavior itself can influence sensory input, we next examined whether sniff magnitude was modulated by perceptual properties of the odor. During the first sniff, both perceived valence (t = 3.1, p = .0017) and intensity (t = 4.6, p = 5e-6) were positively associated with sniff magnitude. During the second sniff, valence remained positively associated with sniff size (t = 6.1, p = 1.4e-9), whereas intensity showed a negative relationship (t = −3.99, p = 6.9e-5), indicating a shift in how odor strength influences sampling across repeated inhalations.

To determine whether second-sniff gamma activity reflects odor valence, spectral responses were compared across pleasant and unpleasant stimuli (Fig. 3). The increase in gamma activity during the second sniff was driven primarily by an increase for unpleasant odors. This effect was visible both when responses were examined separately for pleasant and unpleasant stimuli (Fig. 3A,B,E,F) and when contrasting odor valence categories directly.

**Figure 3:**
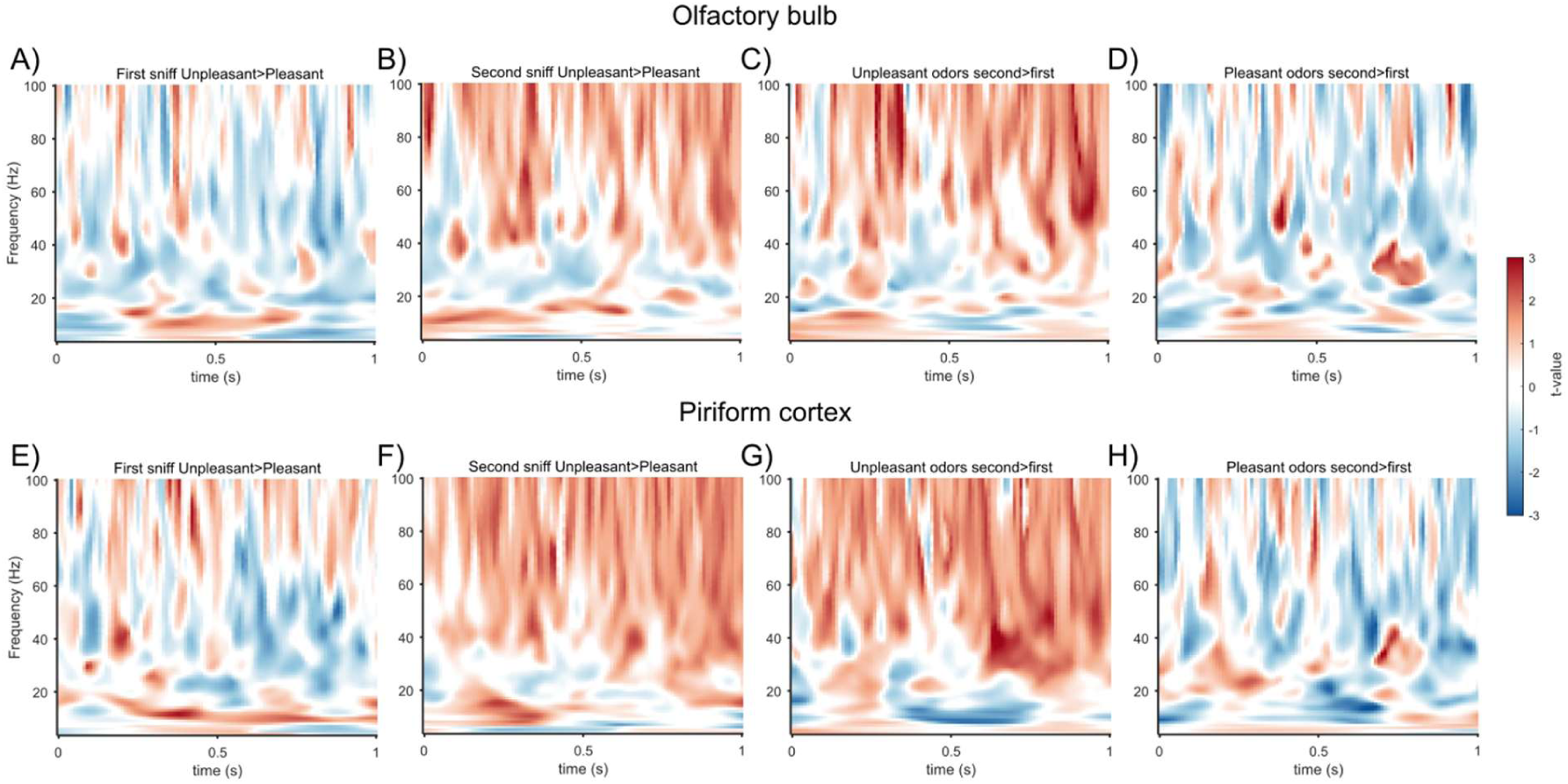
Time-frequency responses contrasting unpleasant and pleasant trials over the first second of the two sniffs show that the stronger gamma activity during the second sniff seems to be driven by unpleasant odor stimuli. **A)** Unpleasantly perceived odors contrasted with pleasantly perceived odors during the first sniff in the OB. **B)** Same as **A,** but for the second sniff. **C)** The second sniff contrasted with the first sniff for unpleasantly perceived odors in the olfactory bulb. **D)** Same as **C,** but for pleasantly perceived odors. **E)**, **F), G), H)** Same as **A, B, C, D** but for the piriform cortex.

A mixed-effects model applied to the full time-frequency spectrum confirmed these findings (Fig. 4), while also controlling for variations in sniffing and in odor intensity. Gamma power increased as a function of negative valence prior to the second sniff and cluster-based permutation testing confirmed a significant valence-informative gamma cluster preceding the second sniff (t=3.4, p=.0013, CI=[.001, .0037], cluster corrected p=.015). The gamma-valence relationship further strengthened during the second sniff, peaking near odor offset (t=4.1, p=.00016, CI=[.0015, .0042], cluster corrected p = .001).

**Figure 4:**
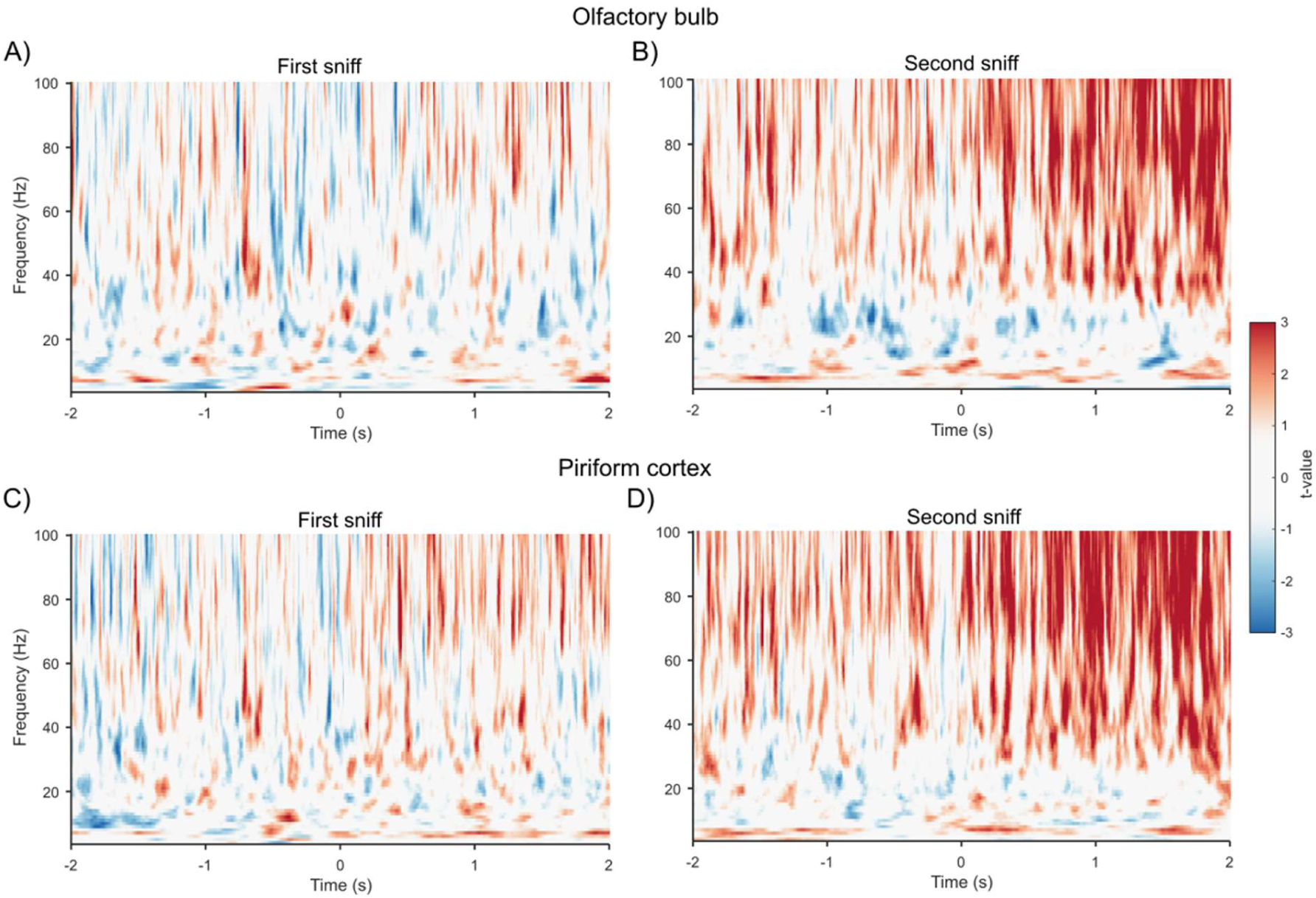
Gamma power after first inhalation is related to odor valence. **A)** Mixed-effects model applied at each time-frequency point of the power spectrum in the OB, predicting spectral power based on perceived valence on a trial-by-trial basis. The perceived valence scale is inverted so that higher negative valence correlates positively with **i**ncreased power. A relationship between odor valence and gamma-band activity is mainly seen 2 seconds after the odor enters the nose. **B)** Same as **A,** but for the second sniff, where a stronger relationship is observed between increased spectral power and perceived negative valence **C)** Same as **A,** but for the PC, showing the same pattern as the OB. **D)** Same as **C,** but for the second sniff in the PC, following the same pattern as the OB.

Additional insights, beyond those offered by traditional wave-based analyses, can be added by analyzing intermittent burst activity because it considers that oscillatory activity is intermittent and varies from trial to trial. Oscillatory bursts are closely linked to population spiking, making them a useful proxy for ensemble activity. We found that gamma burst occurred approximately 0.1 s prior to the onset of the second inhalation (t = 3.9 in OB; t = 3.7 in PC), suggesting a preparedness response.

Together, these findings demonstrate that successive sniffs are accompanied by distinct patterns of oscillatory activity. Whereas the first sniff is characterized by stronger theta-, alpha- and beta-band responses, the second sniff is marked by enhanced gamma activity that provides predictive information about odor valence already before the second sniff. This valence-specific signal intensifies across the inhalation, parallelling the overall rise in gamma power over the two sniffs.

### First-sniff alpha/beta activity predicts second-sniff gamma responses in the olfactory bulb

To determine whether neural activity during one sniff predicts activity during the next, we examined the relationship between first-sniff alpha/beta activity and second-sniff gamma responses in the olfactory bulb.

A linear mixed-effects model with second-sniff gamma as the response variable and first-sniff alpha/beta as a fixed effect revealed a significant association (Fig. 5B), which remained after controlling for pre-stimulus activity (Supplementary Table 1). Notably, the relationship between first-sniff alpha/beta and second-sniff gamma (t = 5.9, p =4.7e-9, CI = [0.070,0.14]) was stronger than that observed for pre-stimulus alpha–beta (t = 0.28, p = .78, CI = [−0.028,0.038]) and pre-stimulus gamma (t = .046, p = .65, CI = [−0.026,0.042]), indicating that second-sniff gamma responses are more closely related to odor-evoked alpha–beta activity than to baseline fluctuations.

**Figure 5:**
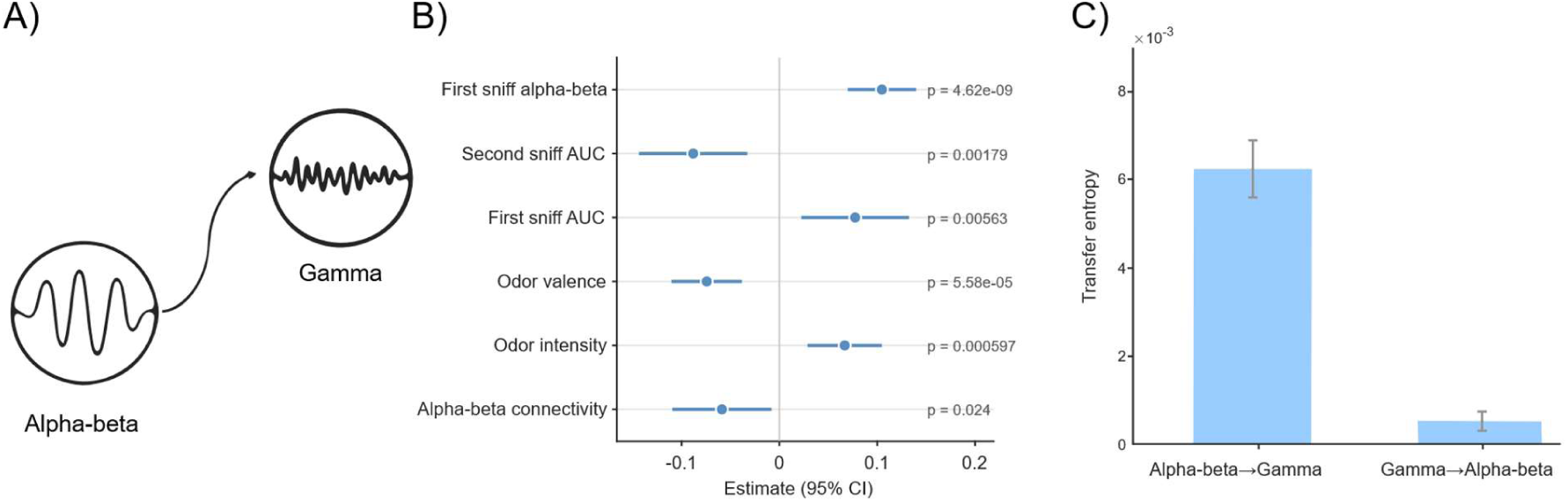
Alpha/beta activity during the first sniff updates gamma-band activity during the second sniff. **A)** Illustration of alpha/beta and gamma activity. **B)** Linear mixed-effects model shows that first sniff alpha/beta predicts second sniff gamma strength. Perceived valence and intensity correlate significantly with second sniff gamma. Further, coherence between the olfactory bulb and piriform cortex during sniff 1 and the size (AUC: area under the curve) of the two sniffs had significant predictive power on the second sniff gamma activity. A table of all the fixed effects can be found in Supplementary Table 1. **C)** Transfer-entropy analysis indicated a causal transfer of information between the alpha/beta band and the gamma band during the first and second sniff through transfer entropy.

Perceptual and network-level factors further contributed to this effect. Both odor valence and perceived intensity predicted second-sniff gamma power, with negative valence and higher intensity associated with stronger gamma responses (Supplementary Table 1). In addition, coherence in the alpha/beta band between OB and PC during the first sniff significantly predicted second-sniff gamma power (t=−2.3, p=.024, CI=[−0.11,-0.007]). Moreover, the size of both the first and the second sniff had a significant influence on the gamma activity of the second sniff, further confirming how integral the sniff is in odor processing.

To assess whether the relationship between the alpha/beta and gamma activity was directional, transfer entropy was estimated across the power spectrum of the OB over the two sniffs (Fig. 5C). Alpha/beta activity predicted subsequent gamma activity significantly more than the reverse pattern (t = 6.5, p <.00001, CI = [.0087,.0165]), indicating a directional flow of information from first-sniff alpha/beta to second-sniff gamma.

Together, these findings demonstrate that second-sniff gamma activity is systematically linked to both alpha/beta activity within the OB and OB-PC functional connectivity during the preceding sniff. This relationship remained after accounting for baseline activity, perceptual ratings, and respiratory parameters, indicating that oscillatory dynamics are linked across successive inhalations rather than reflecting independent sensory responses.

### Piriform cortex alpha/beta activity has stronger functional connectivity to higher order brain regions during the first sniff

We have previously shown that the PC communicates back a richer representation of the odor percept to the OB in the alpha/beta-band during the first sniff. This representation was hypothesized to include coordination of higher-order evaluative networks. To test whether the first sniff preferentially engages higher-order evaluative networks, we examined whole-brain functional connectivity in the alpha–beta range (12 Hz), where power peaked during the first sniff (Fig. 2B and D). We contrasted connectivity between the first and second sniffs to determine whether network interactions were selectively enhanced during initial sampling.

To test whether the first sniff preferentially engages higher-order evaluative networks, we examined whole-brain functional connectivity in the alpha–beta range (12 Hz), where power peaked during the first sniff (Fig. 2B and D). We contrasted connectivity between the first and second sniffs to determine whether network interactions were selectively enhanced during initial sampling.

Connectivity was significantly stronger during the first sniff relative to the second (Fig. 6). The piriform cortex showed its strongest connections with the orbitofrontal cortex (OFC) and insula (Supplementary Table 2). Additional robust connectivity was observed with frontal regions implicated in working memory and with the hippocampus. This pattern indicates that the first sniff engages a distributed evaluation network, whose output may be fed back to influence subsequent neural responses.

**Figure 6:**
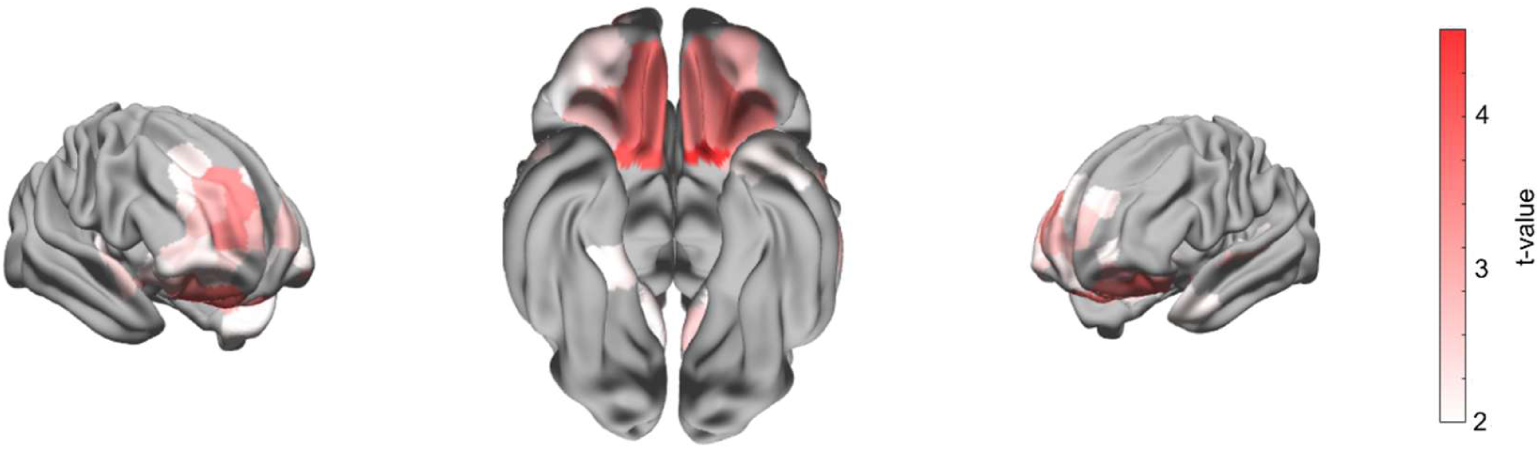
The piriform cortex exhibits stronger functional connectivity with prefrontal and evaluative regions during the first sniff than during the second. These regions included the orbitofrontal cortex, insula, prefrontal areas and the hippocampus.

Network-level metrics further supported this interpretation with frontal regions showing higher node degree during the first sniff (Supplementary Table 2), consistent with their role as connectivity hubs in this period.

## Discussion

Understanding how the human olfactory system maintains perceptual continuity across temporal gaps between inhalations is central to models of olfaction. Yet, the area has received little empirical investigation. Here, we demonstrate that successive odor samples are not processed in isolation but are linked through structured oscillatory interactions between the olfactory bulb (OB) and the piriform cortex (PC). Specifically, first-sniff alpha/beta activity in the OB and coherence between the OB and PC predicted second-sniff gamma power. Transfer entropy analyses supported a directional dependence from first-sniff alpha/beta to second-sniff gamma activity. This cross-sniff interaction persisted after controlling for pre-stimulus oscillatory power and respiratory parameters, suggesting that the network state established during one inhalation shapes processing during the next.

These findings extend previous evidence that beta-and gamma band interactions coordinate OB-PC processing of odor valence (Nordén et al., 2024) and intensity (Nordén et al., 2026). Whereas our earlier work characterized recurrent interactions within a single sniff, the present findings suggest that these interactions also persist across successive inhalations.

We previously suggested that an odor percept emerges through coordinated bottom-up/top-down interactions where the PC returns a refined alpha/beta signal to the OB approximately 0.5–1 s after odor onset (Iravani et al., 2021; Nordén et al., 2024, 2026). Our findings here suggest that this top-down feedback is informed by a network of higher order brain regions, relayed through the PC to update the OB’s local state, priming its gamma output for the subsequent inhalation. The OB thereby functions not merely as a sensory relay, but as a dynamic computational node whose gamma output reflects both afferent receptor input and accumulated top-down influences. This architecture parallels recurrent beta-gamma interactions in visual and auditory cortices that support iterative refinement of sensory representations (Arnal & Giraud, 2012; Michalareas et al., 2016).

This multi-sniff loop is broadly consistent with predictive coding architectures, in which feedforward signals are iteratively refined by slower feedback from higher-order regions (Arnal & Giraud, 2012; Bastos et al., 2012; Keller & Mrsic-Flogel, 2018), as recently proposed for the OB-PC-OFC hierarchy by (Lyons & Gottfried, 2025). However, because our design did not manipulate expectations or include expectation-violation conditions, we cannot definitively establish whether the observed alpha/beta signal carries explicit prediction errors or more generalized evaluative feedback. Our data are equally compatible with a recurrent updating framework in which beta feedback conveys contextual or affective information without computing explicit prediction errors. Alpha/beta activity could also transiently reflect a top-down-dominated network state during the first sniff, while simultaneously setting the conditions under which the OB responds to the next sensory sample. Adjudicating between these accounts will require future designs that parametrically manipulate olfactory expectations across successive sniffs.

The recurrent interactions described above were not independent of perception but were systematically related to subjective odor evaluation. The first sniff was characterized by stronger functional connectivity between the piriform cortex and higher-order evaluative regions, including the orbitofrontal and insular cortices, suggesting that the initial inhalation prioritizes rapid affective assessment and establishes a contextual scaffold for subsequent processing (Gottfried, 2010; Gottfried et al., 2003; Zelano et al., 2011). Gamma-band activity during the second sniff relates to perceived odor valence, with a valence-related gamma increase appearing approximately 100 ms before the onset of the second inhalation and further strengthening throughout the second sniff. This valence specificity was driven primarily by an increase with unpleasant odors, consistent with a prioritized processing of stimuli with high behavioral relevance, where odor valence typically guides approach-avoidance decisions before fine-grained perceptual discrimination is required (Iravani et al., 2021). This temporal dissociation, where the first sniff engages broad evaluative networks and the second sniff carries refined perceptual content in the gamma band, supports the view that successive sniffs serve dissociable computational roles within a single odor encounter (Uchida & Mainen, 2003). One potential concern is that sniff-related differences in airflow could drive the observed spectral changes. However, sniff magnitude and the area under the breathing curve were included as covariates in all mixed-effects models, and the beta-to-gamma relationship survived these controls, indicating that the oscillatory dynamics we report are not reducible to respiratory mechanics.

Much of what is known about the temporal organization of olfactory processing derives from rodent models where odor responses are tightly coupled to respiration-linked theta rhythms that organize beta- and gamma-band activity across sniff cycles (Fontanini & Bower, 2006; Kay & Stopfer, 2006). Recent evidence suggests that a similar principle operates in humans. Using direct recordings from the human olfactory bulb (Sheriff et al., 2026) demonstrated sniff-aligned theta oscillations that coordinate odor-related gamma activity within individual inhalations. Together, these findings suggest that theta-mediated temporal organization may represent a conserved feature of mammalian olfactory processing despite pronounced differences in sampling behavior across species. However, humans sample odors at much lower frequencies (<1 Hz), introducing multi-second gaps between sensory inputs. Our results demonstrate that while intra-sniff theta structures processing within an inhalation, inter-sniff alpha/beta-to-gamma interactions bridge information across inhalations. This multi-timescale organization highlights how human olfaction adapts to sparse sensory sampling by utilizing higher-order cortical networks (OFC, insula, PFC) to maintain network state across physical gaps in input. Together, these findings suggest that human olfactory processing is organized across multiple temporal scales, with theta rhythms structuring neural activity within individual inhalations and recurrent alpha/beta/gamma interactions linking successive odor samples.

Several limitations of the present work warrant consideration. Our design employed only two consecutive sniffs, which leaves open the question whether the beta-to-gamma updating mechanism accumulates across longer sampling sequences or reaches a ceiling after an initial update. The eLORETA-based source reconstruction, while validated for EBG signals (Nordén et al., 2025), imposes spatial resolution constraints that limit our ability to dissociate subregions within the OB or PC. We mitigated this by using predefined ROI coordinates based on stereotactic atlases and by restricting connectivity analyses to broad regional contrasts rather than fine-grained spatial maps. Our stimulus set comprised only four odors varying primarily in valence, and whether inter-sniff updating operates similarly for other perceptual dimensions, such as odor quality or source identity, remains to be tested. Finally, although alpha/beta and gamma activity covaried with perception and predicted subsequent neural responses, these findings do not establish that these oscillations encode information or implement formal updating operations. Rather, they may reflect changes in network state driven by sensory, cognitive, or neuromodulatory influences.

Our findings demonstrate that the human OB-PC circuit does not process each inhalation in isolation but instead carries forward prior perceptual information through alpha/beta-to-gamma interactions that update sensory representations across sniffs. This updating mechanism reconceptualizes the OB as a site of active, history-dependent activity within a recurrent cortical network, where each gamma response is shaped not only by the current afferent input but also by the accumulated evaluative and contextual activity from preceding samples. For humans, whose olfactory system is characterized by seconds of absent sensory input between successive inhalations, such a mechanism is essential for constructing and maintaining a stable odor percept. The subjective continuity of olfactory experience, the sense that an odor is simply “there” despite arriving in discrete packets, may ultimately rest on this capacity of the OB-PC circuit to bridge perceptual gaps through recurrent oscillatory updating. In other words, what we perceive as a single olfactory experience is thus not the product of any one inhalation, but rather the cumulative outcome of a dynamic, iteratively refined neural representation built across successive sniffs.

## Materials and methods

### Participants

Fifty-eight participants were initially recruited through Karolinska Institutet’s online recruitment platform. All participants reported normal olfactory function, no history of neurological or psychiatric disease, and no current nasal congestion or upper respiratory infection. Normal olfactory function was verified using a five-item Sniffin’ Sticks cued-odor identification test (Burghart Messtechnik, Germany). Participants scoring below 3 on the odor ID test or having excessive artifact rejection (>50% of trials in any condition; see Preprocessing) were excluded, yielding a final sample of 48 participants (mean age = 30 ± 8.8 years; 27 women). All participants provided written informed consent prior to participation. The study was approved by the Swedish Ethical Review Authority (Dnr 2017/2332-31/1) and conducted in accordance with the Declaration of Helsinki.

### Odor stimuli and delivery

We employed four odors (Figure 1C), two pleasant: ethyl butyrate (0.25% volume in volume dilution [v/v], Sigma Aldrich, CAS 105-54-4), carvone (50%, Merck, CAS 6485-40-1) and two unpleasant: diethyl disulfide (0.25%, Sigma Aldrich, CAS 110-81-6), fish odor mixture (50%, Symrise Inc.). Their perceived iso-intensity was verified in a pilot study. Odors were delivered birhinally at 3 L/min using a computer-controlled olfactometer (Lundström et al., 2010) with a continuous stream of clean air (0.5 L/min) to minimize tactile cues. The system was flushed with clean air between trials and all air was passed through activated carbon and microfilters to eliminate trace contaminants.

Odor delivery was triggered using E-Prime 2.0 (Psychology Software Tools, Pennsylvania) . The olfactometer has an onset delay of ∼200 ms, the time required for odorant transport through the tubing, which we quantified using a photo-ionization detector (200B miniPID, Aurora Scientific, Ontario). All analyses accounted for this delay, i.e., time 0 in figures corresponds to odor arrival at the nostrils.

### Breathing measurement

Respiration was continuously assessed via a temperature probe placed inside the nostril (MLT415/D, AD Instruments). Decreases and increases in air temperature indexed inhalation and exhalation, respectively. Respiration was sampled at 400 Hz (PowerLab 16/35, ADInstruments, Colorado) and processed in LabChart Pro v7.3.8.

### Procedure

Participants sat in a ventilated, sound-attenuated booth and wore earplugs and headphones playing white noise to mask auditory cues from the olfactometer. To increase signal-to-noise ratio in the EBG measure (Iravani et al., 2020, 2024), participants fasted for six hours before testing. To prevent onset expectancy, all trials were triggered by participants’ sniffs, with inhalation initiating odor delivery. Participants were instructed to breathe normally through their nose throughout the experiment and the triggering mechanism was not disclosed to them.

Each trial began with a fixation cross. After 1–2 s, an odor was delivered at the nadir of inhalation. The odor valve remained open until 1 s after the subsequent inhalation nadir, after which participants rated the odor on visual analog scales for intensity (0–10) and pleasantness (–5 to +5) (Figure 1A).

For analyses requiring a binary pleasant/unpleasant factor, we applied a within-participant median split to subjective pleasantness ratings, independent of stimulus identity. Pleasant trials were rated on average as 6.9 ± .21, and unpleasant trials as 2.0 ± 0.18, *t*(45) = 18.97, *p* = 1.13e-23; Figure 1D.

### EEG and electrobulbogram recording

We recorded neural activity at 512 Hz with 64 scalp electrodes and 4 electrobulbogram electrodes (Figure 1B) using the ActiveTwo system (BioSemi, Amsterdam, The Netherlands). Electrode locations were digitized using an optical neuronavigation system (Brainsight, Rouge Research, Montreal, Canada) and co-registered to standard MNI space. These coordinates were later used for eLORETA source localization.

During acquisition, the signal was high-pass filtered at 0.1 Hz and low-pass filtered at 100 Hz in ActiView (BioSemi). Before recording, all electrode signals were visually inspected, and any electrode with an offset exceeding 40 μV was adjusted until it met the acceptable threshold.

### Signal processing and analysis

#### Preprocessing

We divided the EEG/EBG data into epoch windows spanning 5 s before odor onset and 5 s after odor offset. Because trials were sniff-triggered, trial length varied with participants’ natural breathing. The signals were then re-referenced to the common average, and 50 Hz line noise was removed using a discrete Fourier transform filter. We removed trials with very large muscle artifacts, defined as spectral z-scores ≥ 25 in the 110–140 Hz band and eye blinks using Independent Component Analysis (ICA; InfoMax). The remaining muscle and ocular components were removed using ICA within the SASICA framework (Chaumon et al., 2015), employing the FASTER and ADJUST algorithms. Participants with >50% rejected trials in any odor condition were excluded, yielding 48 participants with an average of 67 and standard deviation of 9 trials.

#### Source time-course reconstruction

To reconstruct the source activity from the OB and PC, we co-registered digitized electrode locations to MNI space using a six-parameter affine transformation and computed the forward model with a Finite Element Method based on the MNI152 template. We segmented the head model into five tissue types (CSF, gray matter, white matter, scalp, skull) using established conductivities [1.79, 0.33, 0.14, 0.43, 0.01] (Vorwerk et al., 2014). A Freesurfer-derived cortical mesh (icosahedral resolution 7) served as the source model and the inverse problem was solved with eLORETA using a 10% regularization parameter. We projected the resulting source time-courses to the principal axis using singular value decomposition. This configuration has shown reliable EBG source estimates (Nordén et al., 2025).

Analyses focused on four predefined ROIs based on stereotactic coordinates: left (−4, 40, −30) and right (4, 40, −30) OB, and left (−22, 0, −14) and right (22, 2, −12) PC. We extracted OB and PC time-courses using identical procedures for all participants. All source reconstruction was performed in FieldTrip 2022 (Oostenveld et al., 2011) in MATLAB 2022a.

#### Power spectrum analysis

We estimated OB–PC spectral power between 4–100 Hz using a multi-taper convolutional method with Hanning tapers and a variable time window that decreased with frequency, ensuring ≥2 cycles per window while maintaining high time resolution. We applied frequency smoothing at 60% of the target frequency to balance temporal precision and spectral resolution across bands. This was chosen because the exact gamma frequency varies between individuals.

To characterize changes in OB and PC dynamics across successive sniffs (Fig. 1A), we performed an analysis in which the spectral power was binned based on the sniff cycle with 1000 bins for the two sniffs. Power spectra were computed on a trial-by-trial basis across both sniffs, resulting in trials of varying duration due to individual differences in breathing cycle length (Supplementary Figure S1).

Statistical significance was evaluated using a 10,000 permutations Monte Carlo simulation with maximum sum cluster-correction.

#### Burst analysis

To characterize oscillatory bursts associated with odor processing, we computed time– frequency representations of the EEG/EBG signal during odor perception. Bursts were defined as transient increases in power exceeding 1 standard deviation above the 10-trial mean and lasting for at least three cycles of the band-specific average frequency (Lundqvist et al., 2016). Because bursts are intermittent and trial-specific, events were detected on single trials within a window extending from 2000 ms before to 2000 ms after the onset of the second sniff.

For each participant, we quantified burst rates separately for the OB and PC in the alpha/beta (8–30 Hz) and gamma (30–100 Hz) bands. To examine how burst activity related to perceived odor valence, we retained single-trial data and fitted linear mixed-effects models for beta and gamma bursts in each region. Odor intensity and sniff size were included as covariates, and participant identity was modeled as a random intercept to account for inter-individual variability. To meet normality assumptions, we applied an inverse-rank transformation to power estimates and verified residual normality using Shapiro–Wilk tests.

#### Linear modelling of the power spectrum

To assess how perceived odor valence predicted neural activity, we quantified time–frequency power using the previously computed power spectra and fitted a linear mixed-effects model at each time–frequency point. Power served as the dependent variable. Fixed effects included perceptual ratings of valence and intensity as well as the area under the curve (AUC) of the breathing trace. To highlight the expected effect, valence ratings were inverted so that positive coefficients correspond to stronger responses for *odors with negative valence*. A random intercept was included for each participant to account for inter-individual variability. To satisfy normality assumptions, we applied an inverse-rank transformation to the power values and verified residual normality using the Shapiro–Wilk test.

Statistical significance was evaluated using a 1,000 permutations Monte-Carlo simulation with maximum sum cluster-correction. We limited the number of permutations to 1,000 due to the exceedingly large computational requirement performing statistical evaluation of parallelized time-frequency spectrums.

#### Linear modelling of the relationship between beta and gamma

To quantify whether alpha/beta activity during the first sniff predicted gamma responses during the second sniff, we extracted alpha/beta-band power (8–30 Hz) from 0.5–1 s after odor onset of the first sniff and gamma-band power (30–100 Hz) during the first second of the second sniff. For each trial, power was averaged across the respective time windows prior to modeling. To isolate this relationship from potential confounds, the model included pre-stimulus beta and gamma activity, perceptual ratings of valence and intensity, and OB–PC coherence as an index of functional connectivity. Coherence was calculated based on the average over multiple tapers per trial. To control for potential respiratory influences, area under the curve (AUC) of the sniff were also entered. A linear mixed-effects model with a random intercept for participant was fit, and the full model specification is provided in Supplementary Equation 1.

Model fit was evaluated against reduced models using Akaike Information Criterion (AIC) and Bayesian Information Criterion (BIC) where the model with the lowest values was selected. To meet normality assumptions, gamma-power values were inverse rank-transformed, and residual normality was verified using Shapiro–Wilk tests.

#### Transfer entropy

To assess directionality, we computed Gaussian Transfer Entropy (TE), an information-theoretic measure of directed statistical dependence between two processes (Schreiber, 2000; Vicente et al., 2011). Positive TE values indicate that the past state of one process improves prediction of the future state of another beyond what can be predicted from its own past. TE was estimated between the beta-band and gamma-band power time courses using the IDTxl library (Wollstadt et al., 2019) in Python 3.14 with multiple temporal lags. Significance was assessed with a permutation framework and false-discovery-rate (FDR) correction at the within-participant level. Values surviving FDR correction were then compared at the group level using two-tailed Student’s *t*-tests.

#### Whole brain connectivity

To evaluate broader network effects, we constructed a whole-brain source model using the Glasser atlas (Glasser et al., 2016), which includes both cortical and subcortical regions. Source reconstruction followed the same pipeline described under *Source time-course reconstruction*, but with 360 dipoles placed at region centroids. Functional connectivity was quantified using cross-spectral density and contrasted between first and second sniff epochs with a Student’s *t*-test.

To characterize network organization, we computed node degree and betweenness centrality and compared these metrics across sniffs. Network analyses were conducted using the Brain Connectivity Toolbox implemented within FieldTrip, and group-level contrasts were performed using two-tailed Student’s *t*-tests.

## Data availability

All anonymized data and scripts required to reproduce the results will be available at zenodo.

## Acknowledgements

This work was supported by the Knut and Alice Wallenberg Foundation (KAW 2018.0152); the ERC Synergy grant D2Smell (Digitizing Smell: From Natural Statistics of Olfactory Perceptual Space to Digital Transmission of Odors; 101118977); and Riksbankens Jubileumsfond (P24-0795).

## Supplementary materials

**Supplementary figure 1:**
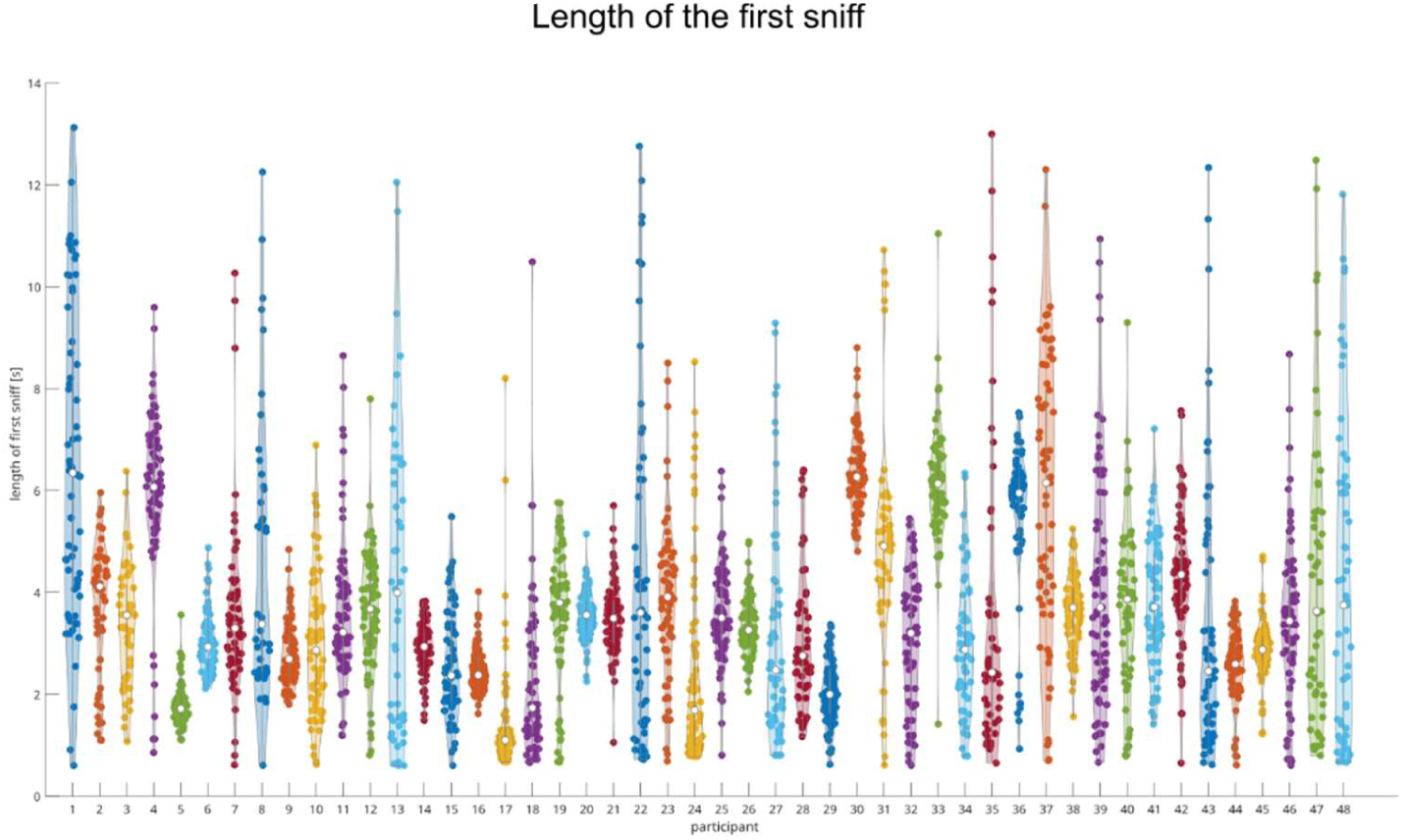
**Sniff length of all participants.**

**Supplementary Table 1:** Linear mixed effect model of first-second sniff dynamics where first sniff alpha/beta predicts second sniff gamma strength.

| Name | t-value | p-value | Lower | Upper |
| --- | --- | --- | --- | --- |
| Intercept | 0.096 | 0.924 | -0.07 | 0.077 |
| First sniff alpha/beta | 5.873 | 4.71e-9 | 0.07 | 0.14 |
| Valence | -4.035 | 5.58e-5 | -0.11 | -0.038 |
| Intensity | 3.436 | 0.001 | 0.029 | 0.105 |
| Pre-stimulus alpha/beta | 0.275 | 0.783 | -0.029 | 0.038 |
| Pre-stimulus gamma | 0.458 | 0.647 | -0.026 | 0.042 |
| Sniff 1 AUC | 2.771 | 0.006 | 0.023 | 0.133 |
| Sniff 2 AUC | -3.126 | 0.002 | -0.144 | -0.033 |
| Sniff 1 alpha/beta coherence | -2.254 | 0.024 | -0.107 | -0.007 |

**Supplementary Table 2: Brain areas with significant connectivity to piriform cortex in the contrast between the first and the second sniff. (See separate file)**

