## Supplementary Table 3 for "Recurrent interactions between olfactory bulb and olfactory cortex support cross-sniff perceptual continuity in humans"

| regionLongName | regionIdLabel | LR | region | Lobe | T_value | regionID |
|---|---|---|---|---|---|---|
| RetroSplenial_Complex_L | 14_L | L | RSC | Par | 2.268668253 | 14 |
| Parieto-Occipital_Sulcus_Area_1_L | 31_L | L | POS1 | Par | 2.086536814 | 31 |
| Area_ventral_23_a+b_L | 33_L | L | v23ab | Par | 2.493434251 | 33 |
| Area_a24_L | 61_L | L | a24 | Fr | 2.799481643 | 61 |
| Area_dorsal_32_L | 62_L | L | d32 | Fr | 2.040433621 | 62 |
| Area_p32_L | 64_L | L | p32 | Fr | 2.677366254 | 64 |
| Area_10r_L | 65_L | L | 10r | Fr | 2.801571075 | 65 |
| Area_47m_L | 66_L | L | 47m | Fr | 2.739099894 | 66 |
| Area_8Ad_L | 68_L | L | 8Ad | Fr | 2.214406718 | 68 |
| Area_9_Middle_L | 69_L | L | 9m | Fr | 2.090682765 | 69 |
| Area_9_Posterior_L | 71_L | L | 9p | Fr | 3.315624614 | 71 |
| Area_10d_L | 72_L | L | 10d | Fr | 2.752800986 | 72 |
| Area_47l_(47_lateral)_L | 76_L | L | 47l | Fr | 2.020065211 | 76 |
| Area_anterior_47r_L | 77_L | L | a47r | Fr | 2.183004329 | 77 |
| Area_46_L | 84_L | L | 46 | Fr | 2.226452231 | 84 |
| Area_anterior_9-46v_L | 85_L | L | a9-46v | Fr | 2.630896458 | 85 |
| Area_9-46d_L | 86_L | L | 9-46d | Fr | 2.870300982 | 86 |
| Area_9_anterior_L | 87_L | L | 9a | Fr | 3.329945999 | 87 |
| Area_10v_L | 88_L | L | 10v | Fr | 2.200680538 | 88 |
| Area_anterior_10p_L | 89_L | L | a10p | Fr | 2.461093969 | 89 |
| Area_11l_L | 91_L | L | 11l | Fr | 2.356182231 | 91 |
| Area_13l_L | 92_L | L | 13l | Fr | 3.317332738 | 92 |
| Orbital_Frontal_Complex_L | 93_L | L | OFC | Fr | 3.648782192 | 93 |
| Area_47s_L | 94_L | L | 47s | Fr | 2.666619324 | 94 |
| Area_52_L | 103_L | L | 52 | Temp | 2.169739687 | 103 |
| Area_TA2_L | 107_L | L | TA2 | Temp | 2.457095455 | 107 |
| Pirform_Cortex_L | 110_L | L | Pir | Temp | 2.968124584 | 110 |
| Anterior_Ventral_Insular_Area_L | 111_L | L | AVI | Fr | 2.002759168 | 111 |
| Anterior_Agranular_Insula_Complex_L | 112_L | L | AAIC | Ins | 2.4101196 | 112 |
| PreSubiculum_L | 119_L | L | PreS | Temp | 2.042649736 | 119 |
| Hippocampus_L | 120_L | L | H | Temp | 2.22514712 | 120 |
| Area_STGa_L | 123_L | L | STGa | Temp | 2.451935543 | 123 |
| ParaHippocampal_Area_1_L | 126_L | L | PHA1 | Temp | 2.202311211 | 126 |
| ParaHippocampal_Area_2_L | 155_L | L | PHA2 | Temp | 2.065508526 | 155 |
| Area_25_L | 164_L | L | 25 | Fr | 3.290471936 | 164 |
| Area_s32_L | 165_L | L | s32 | Fr | 3.28483201 | 165 |
| Posterior_OFC_Complex_L | 166_L | L | pOFC | Fr | 3.963993045 | 166 |
| Area_Posterior_Insular_1_L | 167_L | L | PoI1 | Ins | 2.266602957 | 167 |
| Area_posterior_10p_L | 170_L | L | p10p | Fr | 3.34173978 | 170 |
| Area_posterior_47r_L | 171_L | L | p47r | Fr | 2.34683162 | 171 |
| Para-Insular_Area_L | 178_L | L | PI | Temp | 3.655904904 | 178 |
| Area_posterior_24_L | 180_L | L | p24 | Fr | 2.229574647 | 180 |
| Parieto-Occipital_Sulcus_Area_1_R | 231_R | R | POS1 | Par | 2.619323466 | 231 |
| Area_ventral_23_a+b_R | 233_R | R | v23ab | Par | 2.294820046 | 233 |
| Area_a24_R | 261_R | R | a24 | Fr | 2.00513672 | 261 |
| Area_10r_R | 265_R | R | 10r | Fr | 2.099575392 | 265 |

| | | | | | | |
|---|---|---|---|---|---|---|
| Area_47m_R | 266_R | R | 47m | Fr | 3.210907077 | 266 |
| Area_10d_R | 272_R | R | 10d | Fr | 2.588337323 | 272 |
| Area_9_anterior_R | 287_R | R | 9a | Fr | 2.258344228 | 287 |
| Area_anterior_10p_R | 289_R | R | a10p | Fr | 2.127861549 | 289 |
| Area_11l_R | 291_R | R | 11l | Fr | 2.896067856 | 291 |
| Area_13l_R | 292_R | R | 13l | Fr | 3.695452601 | 292 |
| Orbital_Frontal_Complex_R | 293_R | R | OFC | Fr | 3.438842922 | 293 |
| Area_47s_R | 294_R | R | 47s | Fr | 3.402134095 | 294 |
| Area_OP2-3-VS_R | 302_R | R | OP2-3 | Par | 2.114334287 | 302 |
| Pirform_Cortex_R | 310_R | R | Pir | Temp | 2.968124584 | 310 |
| Anterior_Ventral_Insular_Area_R | 311_R | R | AVI | Ins | 2.303345271 | 311 |
| ProStriate_Area_R | 321_R | R | ProS | Par | 2.167799828 | 321 |
| ParaBelt_Complex_R | 324_R | R | PBelt | Temp | 2.151578638 | 324 |
| Auditory_5_Complex_R | 325_R | R | A5 | Temp | 2.814646271 | 325 |
| Area_TG_dorsal_R | 331_R | R | TGd | Temp | 2.123810847 | 331 |
| Area_25_R | 364_R | R | 25 | Fr | 2.799033861 | 364 |
| Area_s32_R | 365_R | R | s32 | Fr | 2.776601749 | 365 |
| posterior_OFC_Complex_R | 366_R | R | pOFC | Fr | 4.443206573 | 366 |
| Insular_Granular_Complex_R | 368_R | R | Ig | Ins | 2.111346219 | 368 |
| Para-Insular_Area_R | 378_R | R | PI | Temp | 2.068981331 | 378 |

| Cortex_ID | x_cog | y_cog | z_cog | volmm |
|---|---|---|---|---|
| 18 | 96.918149 | 86.60261 | 88.986951 | 843 |
| 18 | 101.731298 | 67.452417 | 84.880916 | 1965 |
| 18 | 94.704348 | 70.024638 | 91.217391 | 690 |
| 19 | 96.046431 | 166.310464 | 66.976438 | 1443 |
| 19 | 98.638734 | 168.099317 | 96.929236 | 1611 |
| 19 | 101.628302 | 174.318868 | 67.658491 | 530 |
| 19 | 96.816092 | 176.014368 | 59.294061 | 1044 |
| 20 | 127.422402 | 156.510121 | 55.184885 | 741 |
| 22 | 113.687799 | 154.661483 | 115.801037 | 2508 |
| 19 | 97.019809 | 180.108831 | 93.67494 | 4190 |
| 22 | 109.307738 | 173.069108 | 109.713526 | 1693 |
| 20 | 100.894288 | 191.564911 | 78.3023 | 2696 |
| 21 | 136.797521 | 154.852273 | 60.41374 | 1936 |
| 21 | 131.095281 | 174.396953 | 59.360514 | 3348 |
| 22 | 127.033498 | 164.498689 | 105.07457 | 3433 |
| 22 | 131.147188 | 176.872535 | 80.33382 | 2738 |
| 22 | 120.191583 | 172.262563 | 93.433103 | 3184 |
| 22 | 112.60695 | 183.845628 | 95.711763 | 3511 |
| 19 | 95.085146 | 179.625174 | 52.61453 | 2161 |
| 20 | 116.405009 | 185.963731 | 63.137306 | 1158 |
| 20 | 115.577956 | 172.905411 | 55.238477 | 2495 |
| 20 | 113.105234 | 153.630846 | 50.639198 | 1796 |
| 20 | 101.248092 | 155.905125 | 46.772628 | 1834 |
| 20 | 123.285457 | 146.215144 | 51.115986 | 1664 |
| 10 | 130.277162 | 105.017738 | 70.645233 | 451 |
| 11 | 141.08604 | 129.967515 | 63.526778 | 1139 |
| 12 | 122.938706 | 130.870602 | 53.682179 | 881 |
| 12 | 121.113485 | 150.508678 | 67.823765 | 749 |
| 12 | 126.225504 | 139.71902 | 60.595101 | 1388 |
| 13 | 109.856877 | 94.634758 | 60.597584 | 1076 |
| 13 | 120.203438 | 102.589542 | 54.63467 | 1396 |
| 11 | 139.957301 | 140.979968 | 50.18292 | 1897 |
| 13 | 111.705833 | 90.678333 | 55.309167 | 1200 |
| 13 | 121.389831 | 90.169492 | 58.122881 | 236 |
| 19 | 95.291898 | 147.049945 | 57.497225 | 901 |
| 19 | 97.364516 | 159.174194 | 54.480645 | 310 |
| 19 | 104.455746 | 137.165786 | 52.015852 | 1514 |
| 12 | 129.839378 | 113.685665 | 68.664076 | 1158 |
| 20 | 114.061006 | 189.546541 | 72.976101 | 1590 |
| 21 | 134.951128 | 169.179699 | 72.101504 | 1330 |
| 12 | 134.650549 | 124.293407 | 56.102198 | 910 |
| 19 | 95.442887 | 162.526279 | 84.888577 | 1427 |
| 18 | 79.376277 | 69.248778 | 87.420702 | 2251 |
| 18 | 86.05698 | 74.521368 | 90.964387 | 702 |
| 19 | 85.878669 | 164.201566 | 70.600783 | 1533 |
| 19 | 85.146262 | 172.751896 | 60.15493 | 923 |

| | | | | |
|---|---|---|---|---|
| 20 | 57.754386 | 157.371517 | 53.686275 | 969 |
| 20 | 81.086781 | 193.730216 | 74.665468 | 2224 |
| 22 | 70.836969 | 188.89409 | 90.121777 | 2521 |
| 20 | 66.79639 | 188.730686 | 61.363177 | 1385 |
| 20 | 65.791587 | 172.746973 | 54.321224 | 3138 |
| 20 | 70.701739 | 152.646957 | 50.322609 | 1150 |
| 20 | 83.615202 | 158.638955 | 45.933017 | 2105 |
| 20 | 60.445039 | 146.166964 | 51.135472 | 1683 |
| 9 | 51.053528 | 111.892944 | 90.335766 | 411 |
| 12 | 57.555081 | 133.362084 | 52.703672 | 1171 |
| 12 | 58.445161 | 150.95914 | 67.72043 | 930 |
| 18 | 70.101423 | 77.854093 | 73.617438 | 562 |
| 10 | 33.096591 | 107.575487 | 79.568994 | 1232 |
| 11 | 29.134363 | 113.095062 | 68.029224 | 2977 |
| 14 | 55.264395 | 140.071941 | 34.559763 | 7103 |
| 19 | 87.037037 | 143.861454 | 58.698217 | 729 |
| 19 | 86.948307 | 156.390374 | 55.54902 | 561 |
| 19 | 77.656757 | 138.762838 | 52.856757 | 1480 |
| 12 | 54.984127 | 112.0839 | 86.482993 | 441 |
| 12 | 47.013363 | 123.157016 | 57.926503 | 898 |
